# MUMdex: MUM-based structural variation detection

**DOI:** 10.1101/078261

**Authors:** Peter A. Andrews, Ivan Iossifov, Jude Kendall, Steven Marks, Lakshmi Muthuswamy, Zihua Wang, Dan Levy, Michael Wigler

**Affiliations:** Simons Center for Quantitative Biology, Cold Spring Harbor Laboratory, Cold Spring Harbor, NY 11724, USA; New York Genome Center, New York, NY 10013, USA.

## Abstract

**Motivation:** Standard genome sequence alignment tools primarily designed to find one alignment per read have difficulty detecting inversion, translocation and large insertion and deletion (indel) events. Moreover, dedicated split read alignment methods that depend only upon the reference genome may misidentify or find too many potential split read alignments because of reference genome anomalies.

**Methods:** We introduce MUMdex, a Maximal Unique Match (MUM)-based genomic analysis software package consisting of a sequence aligner to the reference genome, a storage-indexing format and analysis software. Discordant reference alignments of MUMs are especially suitable for identifying inversion, translocation and large indel differences in unique regions. Extracted population databases are used as filters for flaws in the reference genome. We describe the concepts underlying MUM-based analysis, the software implementation and its usage.

**Results:** We demonstrate via simulation that the MUMdex aligner and alignment format are able to correctly detect and record genomic events. We characterize alignment performance and output file sizes for human whole genome data and compare to Bowtie 2 and the BAM format. Preliminary results demonstrate the practicality of the analysis approach by detecting *de novo* mutation candidates in human whole genome DNA sequence data from 510 families. We provide a population database of events from these families for use by others.

**Availability:** http://mumdex.com/

**Contact:** (or)

**Supplementary information:** Supplementary data are available online.

## 1 Introduction

Standard genome sequence alignment tools, such as Bowtie 2 (Langmead *et al*., 2012) or BWA (Li, Durbin, 2009a), are primarily designed to find one alignment per read while allowing for soft clipping, base substitutions and small insertions and deletions (indels). Existing analysis software for detecting structural variants typically starts with conventional alignment tools, and then looks for either discordant read pair mates (Chen *et al*., 2009; Lindberg *et al*., 2015) or split read alignments (Karakoc *et al.,* 2012; Ye *et al.,* 2009) with read pair mate support. Discordant read pair analysis methods have trouble identifying precise breakpoints and identifying small indels, while split read methods are limited to smaller indels if the underlying aligner allows for non-unique or mutated alignments, since the number of possibilities for alignment would then be too large.

We introduce MUMdex, a Maximal Unique Match (MUM)-based genomic analysis software package for sequence analysis. A MUM between two sequences is defined as an exact match subsequence that exists only once in each sequence (is unique) and is not part of any longer exact match (is maximal). Finding MUMs is computationally rapid and allows us to find all the MUMs between billions of short sequencing reads and the large human reference genome. Pairs of MUMs within a read that have incompatible reference coordinates are starting points for inference about sequence structure. MUMdex software allows events of any size to be confidently detected.

The MUMdex aligner saves read pair information in an indexed lossless compact binary format as MUMs plus the sequence not covered by MUMs. This format facilitates subsequent searching for genomic rearrangements of all kinds by inspecting each pair of MUMs (called a ‘bridge’) within a read. MUMdex analysis software computes a numerical ‘invariant’ for each bridge. When bridge invariants occur with nonzero values, and are seen in multiple independent reads, they signal either genome rearrangements (inversions, translocations or indels) or problems in the reference genome. By comparing the bridge invariants from cancer to normal from the same individual, or from an individual to its parents or to populations, most errors caused by the imperfect reference genome can be eliminated, thus reliably detecting *de novo* and rare rearrangements of any size. MUMdex analysis software can also detect single nucleotide polymorphisms (SNPs), but it is expected to underperform standard methods for detection in regions of high divergence from the reference genome.

## 2 Methods

The core of our method is the ‘bridge invariant’ that is associated with differences between sample and reference genomes. Imagine any genomic rearrangement in which one unique piece of the genome is joined to another unique piece. Each piece has coordinates in a reference genome, and these coordinates can be extended locally to adjacent base pairs even beyond the join. There are two such coordinate extensions at every base. These coordinates will conflict, but either the difference or the sum of the two coordinates at each base will have the same nonzero value. The event type dictates whether it is the difference or the sum that produces the constant. We call this constant the ‘bridge invariant’.

The invariant can be computed from any sequence read that spans the join with enough sequence to establish MUMs on either side of the join. The properly computed invariant will be independent of the strand of the read, the start position of the read, the read length, or base calling errors. The computation will be dependent on the reference coordinate system and so where the reference assembly reflects a minor variant in the human population there will be an associated nonzero bridge invariant in most individuals. For this and other reasons, we construct a database containing nonzero invariants associated with locations from normal genomes. This database is helpful as a filter for events, whether we seek differences between a cancer and the host normal or between a child and its parents.

### 2.1 MUMs, Anchors, Bridges and Invariants

A read is evidence for the sequence structure of the sampled genome (see Figure 1). Most reads will contain one or more MUMs to the reference genome, aligning either in its 5’ to 3’ orientation (‘forward’) or in its reverse complement (‘reverse’). If a MUM terminates within but not at the edge of a read, the final base of the MUM at that termination point is called an ‘anchor’. Every anchor inherits from its MUM a reference genome coordinate, the ‘anchor position’. An anchor with the lowest reference coordinate in the MUM is called the ‘low’ anchor while one with the highest is called the ‘high’ anchor. The presence of an anchor means that the read is not entirely consistent with the reference genome. The sequence adjacent to an anchor but outside the MUM is either a sequence error or a true difference between the sample and the reference.

**Figure 1.**
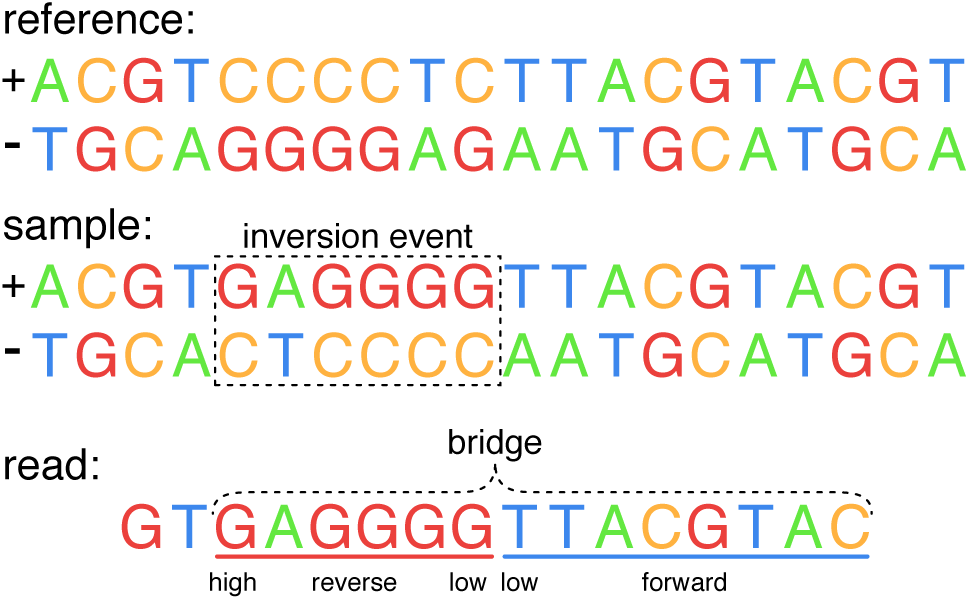
MUMs, bridges and anchors. A ‘reference’ genome, a ‘sample’ genome with an inversion event and a read from the sample are shown. The sequence CCCCTC on the forward (+) strand of the reference genome has been replaced by its reverse complement of GAGGGG in the sample. The read is a subsequence of the sample genome forward strand. The left red underlined MUM and right blue underlined MUM form a bridge, depicted as a bracket on top of the MUMs. Low and high MUM anchor locations and alignment orientations are labeled on the bottom. The right MUM has no high anchor because it terminates at the edge of the read. The low anchors are the bridge anchors for this bridge.

We call any pair of MUMs in a read a ‘bridge’. The MUMs of a bridge may overlap, abut or have a gap between them in the read. The ‘bridge anchors’ are the ‘left bridge anchor’ (the right anchor of the left MUM) and the ‘right bridge anchor’ (the left anchor of the right MUM). If not due to sequence error, bridges arise from substitutions, indels, inversions or translocations, and these types can be distinguished by bridge invariants, as we now discuss.

A bridge can be characterized by the bridge anchor ‘offset’, which is the right bridge anchor read coordinate minus the left bridge anchor read coordinate. The offset is positive if the MUMs do not overlap. It is also characterized by the bridge anchor types (low or high) and the genomic coordinates of the bridge anchors. In the absence of read error, these characteristics together with any sequence separating the MUMs uniquely characterize an event: the characteristics are unaffected by differences in read strand, read length and read placement. For example, the lengths of the bridge MUMs do not affect the bridge characteristics.

However, to obtain a characteristic that is not sensitive to substitution error, and that can be used between individuals who might differ by SNPs, we need something stronger, the bridge invariant. The bridge invariant plays a critical role when we search for structural genomic variation.

The bridge invariant is calculated from the other bridge characteristics, but is not sensitive to substitution read error or substitution polymorphisms. The invariant also is useful for typing an event, and, for simple events, measuring its size.

To calculate the bridge invariant, we begin by picking any base position *b* with a read coordinate *R*_*b*_ within MUM *m*. Based on the alignment of the MUM to the reference genome, *b* has a unique genome coordinate *G*_*m,b*_. Now consider any read position *x* with read coordinate *R*_*x*_. The MUM *m* induces a MUM-genome coordinate for that position, *G*_*m,x*_, namely

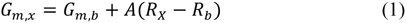

where *A* is +1 for forward MUMs and −1 for reverse MUMs.

Note that *G*_*m,x*_ will be independent of the choice of *b*. Let *n* be the other MUM. Then, up to a sign, the bridge invariant *I* is evaluated at any base *x* as the *difference* of the *m*- and *n*- genome coordinates when *m* and *n* have the *same* orientation (forward or reverse), and the *sum* of the coordinates if they have *different* orientations. To resolve the ambiguity in sign when computing we use this formulation for the bridge invariant:

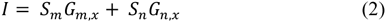
where for each MUM *j*, *S*_*j*_ is −1 for a MUM if it has a low bridge anchor and +1 if it has a high bridge anchor. Therefore, the invariant is 0 if the bridge is caused by base substitutions, negative if caused by deletions and positive if caused by insertions. For indels the absolute value indicates the length of the event. Figure 2 shows an example of an invariant calculation for a small insertion event.

**Figure 2.**
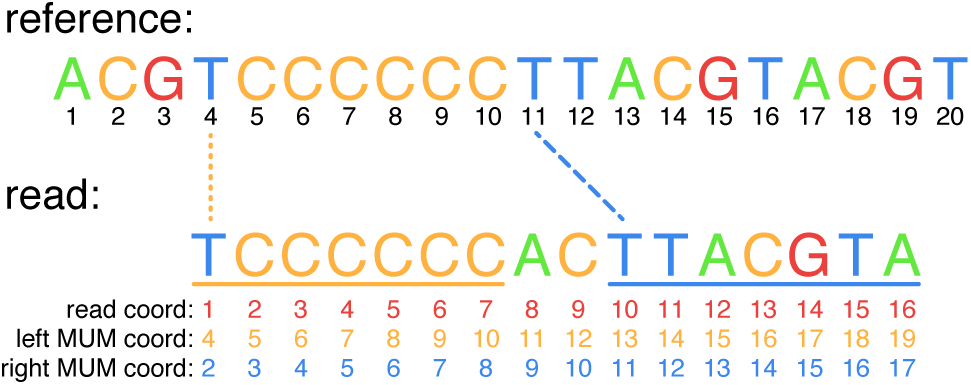
Computing the invariant. A reference genome and an insertion event in a read are shown. Genomic coordinates are displayed in black underneath the reference sequence. The sequence AC has been inserted into the sample genome between positions 10 and 11 of the reference genome. A read from the sample genome is shown, with read coordinates displayed underneath in red. Dotted and dashed lines colored the same as each underlined MUM in the bridge demonstrate the genomic coordinates induced by the MUMs on the read at the first base of each MUM. The coordinates induced on the read by the MUMs are displayed underneath the read in yellow and blue, and note that the difference in the two coordinate systems is a constant. Arbitrarily choosing *x* as the first base of the right MUM (position 10 in the read coordinates) to evaluate the genomic coordinates induced by both MUMs, we note that the left MUM with a high bridge anchor induces coordinate 13 (yellow) and the right MUM with a low bridge anchor induces coordinate 11 (blue), so the bridge invariant is +13 + -11 = 2 according to Equation 2.

#### 2.1.1 Spurious MUMs

Both read error and real events can create novel sequence that by chance matches uniquely to a completely unrelated portion of the genome. We call these chance matches *spurious* MUMs. All aligners encounter spurious alignments but they are typically filtered out on the fly with some degree of success. In contrast, MUMdex retains all MUMs passing optional filters because even spurious MUMs can signal actual events. For example, a MUM, even if spurious, seen recurrently in one child but not in either parent may suggest a real *de novo* event.

To quantify the occurrence of spurious MUMs due to read error, we used the MUMdex aligner (described later in Software) to align 100 million 151 base long simulated reads containing one centered single base substitution between unique sequences from the hg19 human reference genome. In addition to finding the two expected MUMs, we also found on average 6.02 (unfiltered) spurious MUMs per read. With simple filters we do much better. First, by requiring that every MUM be at least 20 bases long, 85.3 percent of spurious MUMs are eliminated. Secondly, we define ‘excess mappability’ as the length of a MUM minus the minimum length required to achieve uniqueness for a subsequence within the MUM. Just requiring MUMs to have an excess mappability of 2 or more eliminates 89.2 percent of spurious MUMs. Additional filters, such as requiring that supporting MUMs be seen on read pair mates can also help to reduce spurious MUM contamination of called events.

#### 2.1.2 Bridges and Invariants for Selected Event Types

Substitutions, deletions and insertions of any length, inversions and translocations can be identified by their distinctive bridge characteristics. A guide to the bridge structures for these types of events is tabulated in Supplementary Figure 1.

Bridges bracketing sequence substitutions have similarly oriented MUMs and a zero invariant. Bridges for deletion events have similarly oriented MUMs but a negative invariant equal to the length of the deleted sequence. These are the simplest events.

A simple tandem duplication may result in a single bridge with an invariant equal to the length of the duplicated sequence. Many more complications can arise from microsatellite expansion and contraction.

A non-tandem insertion event may result in up to three bridges. One bridge spanning the insertion will have similarly oriented MUMs and a positive invariant equal to the inserted sequence length. If the inserted piece contains a MUM, two other bridges are generated: one from the left flank to the inserted sequence; and one from the inserted sequence to the right flank. Those bridge invariants are more like those from translocations, and do not have quantitative interpretation. Of course, if an insertion is too large, the first type of bridge might not be observed. Instead, bridges at the junctions may be observed separately.

Inversion events can produce a distinctive signature: similar absolute value positive and negative invariants located nearby, with absolute value close to twice the local genomic coordinate. If a perfectly inverted sequence (no removal or addition of bases) is short enough to be bracketed by MUMs in a read, the inverted sequence acts as a substitution so the bracketing bridge will have zero invariant. We use an expansive definition of inversion throughout this paper, calling any bridge with two MUMs on the same chromosome but aligning to opposite strands an inversion.

Translocations are defined as bridges with MUMs on different chromosomes that cannot be identified as part of an insertion event. Translocations are expected to be rare, but most bridges resulting from spurious MUMs will be cross-chromosome. Translocations are therefore suspect and may require more evidence, such as increased excess mappability, more read pair mate support, and the absence of the event in the population database.

### 2.2 MUM-Based Analysis

#### 2.2.1 Searching for *de novo* events

To illustrate how we use the processing (storage and analysis) system we have created, we consider a particular application, one of many, the occurrence of a *de novo* structural event. The signature for such an event is clear: we find in the child bridges in multiple reads each at the same location and with the same nonzero invariant, and not in the parents despite high depth of coverage at the same location. The data structures we build are designed to facilitate such searches. We continue to refer to this application in the following.

#### .2.2.2 Recurrence and coverage, operational conditions

Under operating conditions, base-calling errors will be common, coverage may be variable, and peculiarities in the reference genome relative to the child all can cause misinterpretation of data. In this section we discuss these, and explain the auxiliary features of the processing that lessen their impact. We introduce the notions of ambient coverage, anchor counts, support from paired end alignment, and so on. These lead to quantitative filters that can be applied with varying degrees of stringency.

Base calling errors may cause misalignment and result in spurious MUMs and invariants. Fortunately, most read errors are not recurrent, so we can set as a requirement a threshold for number of recurrent invariants. We count invariants by read pair; if an invariant was seen in both mates for a read pair it is counted only once.

Similarly, coverage is important in the parents. If we fail to see the invariant in one of the parents, it could be a failure of coverage or merely the result of under-sampling. Although this is unavoidable on occasion, we take several steps to guard against it. First, we define ‘ambient coverage’ of a genomic position for a sample as how many read pairs in the sample contain a MUM that is at least 25 bases long (to exclude most spurious MUMs) that covers the position. We require that the ambient coverage at both bridge anchor positions of the event in both parents exceed a reasonably large threshold.

Second, we check for isolated bridge anchors in the parents, to see if the adjacent sequence is mostly compatible with the child’s consensus sequence for the event. This allows us to disqualify events as *de novo* if the parent had the event but one of the MUMs is not present due to base calling error, somatic mutation or just being cut off at the end of a read.

Additionally, we search for bridges in a larger population, because observing the same bridge in the general population can be taken as evidence that the event is common, and merely missed in one of the parents. Moreover, bridges that are highly common in certain regions may indicate potential flaws in the reference genome, or else regions that are not stable during library preparation. It is important to note, however, that certain regions of the genome may be highly unstable within the germline or somatic cells of the individual as we shall discuss later.

We also seek to determine that the bridge MUMs in the child themselves are long enough that a single nucleotide polymorphism in the region does not cause recurrent spurious alignment. Along the same lines, we seek in the paired reads evidence that the sequences adjacent to an event are consistent with our interpretation of the event, for example that the right paired read is consistent with the rightmost MUM alignment. We therefore require that each MUM in a bridge has consistent read pair mate support in at least one read pair.

## 3 Software

The MUMdex package consists of an aligner, an alignment format, analysis software and a portable population database of common structural variants to aid filtering. The MUMdex alignment format contains all MUM alignments to the reference found in read pairs, indexes read pairs by the genomic coordinates of their MUMs, and is able to reconstitute input read pair sequences. MUMdex analysis software examines MUMdex alignment files for a population to detect rare and *de novo* mutation candidates.

MUMdex software is written in C++ for the C++14 (ISO/IEC, 2014) standard but is also compilable using C++11. It has been tested most fully using the GCC compiler version 4.9.2 in the Linux operating system. Considerable attention was paid to output format compactness and analysis efficiency. There is also an optional Python wrapper which can run the MUMdex aligner and allows complete access to the MUMdex alignment format.

### 3.1 The MUMdex Aligner

The MUMdex aligner (mummer) uses a suffix array (Manber *et al*., 1993) and Longest Common Prefix (LCP) array (Kasai *et al*., 2001) to efficiently find a variant of a MUM where the sequence may be repeated in the query. This variant (called a maximal almost unique match, or MAM) allows for discovery of tandem duplications in addition to all other mutation types such as SNPs, indels of any size, inversions and translocations.

The MUMdex aligner does not utilize base quality score information generated by the sequencing instrument. A reasonable pre-processing step prior to MUMdex alignment (that we do not perform) would be to clip the ends of reads if quality scores become unacceptable.

The MUMdex aligner borrowed the suffix array implementation of the sparseMEM package (Khan *et al*., 2009) and extensively modified it to:

1. provide object-oriented interfaces and increase parallelism
2. remove sparse feature to boost speed and lower complexity
3. remove a genome length limitation of 2.147 billion bases
4. allow saving and regular or memory-mapped loading of a binary reference and the suffix array and LCP structures
5. read query input in the SAM (Li *et al*., 2009b) or FASTQ (Cock *et al*., 2010) formats
6. optionally pass through quality scores and SAM fields
7. eliminate multiple parsing of query input
8. align to the reference and its reverse complement
9. save sequences and alignments in the MUMdex format

The MUMdex aligner will automatically generate and save the binary reference, suffix array and LCP array if they do not yet exist. The suffix array generation process for a human reference genome requires less than 32 GB of physical memory and may take several hours. Subsequent memory-mapped use of the suffix array and associated structures is capable of performing alignment with only a few GB of physical memory, but will proceed much more efficiently if at least 32 GB of physical memory is available. Optional simultaneous alignment (producing identical output) to the reference and its reverse complement requires 120 GB of memory but is 3x faster.

The MUMdex aligner outputs the MUMdex format directly in a set of separate MUMdex format subdirectories of fixed maximum number of read pairs, pre-sorted to put likely duplicate read pairs adjacent to each other. The program merge_mumdex merges the MUMdex parts into a single MUMdex output directory, marks duplicate read pairs, generates and saves the genome order MUM index and then removes the MUMdex parts (Supplementary Figure 2).

### 3.2 The MUMdex Alignment Format

The MUMdex alignment format stores read pair, MUM and sequence information in memory or in files as arrays of POD (Plain Old Data) C++ objects in native binary format (Figure 3). This means the output format may not be portable between different machine architectures or compilers, but data access can be very fast.

**Figure 3.**
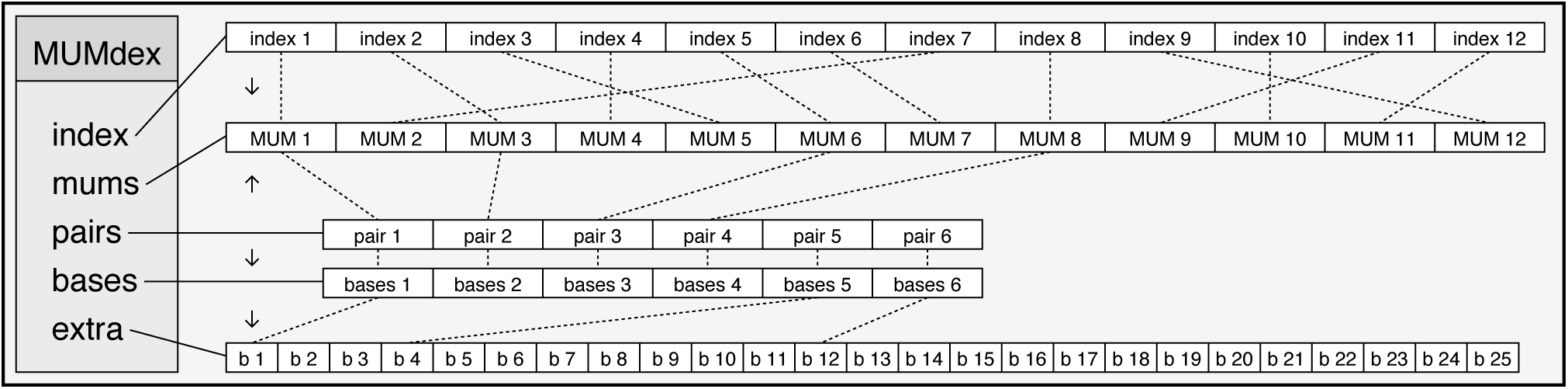
Topology of the MUMdex alignment format. The MUMdex object contains 5 arrays of objects. Arrows at the left show the direction of linkage between objects depicted as dotted lines. The pairs array stores pair objects, sorted by the lowest reference coordinate of the first MUMs in each mate. Pair objects point to their associated bases object and the first MUM in the pair. According to this scheme, pairs 5 and 6 in the diagram have no MUMs while pair 2 has 3 MUMs. MUM index objects point to MUMs via their pairs (full linkage not shown) and are sorted in genome order for the MUMs they point to. The bases array stores one bases object for each pair to encode pair sequence not covered by MUMs. If the sequence is too long to fit in a bases object, the bases object points to a block in the extra array to store the sequence.

All MUMs for a read pair are stored in a block and the MUM blocks and sequence information are stored in the same order as the read pairs. All information for each read pair can therefore be very efficiently streamed into memory (using memory-mapped files). An index enables optional traversal of all MUMs and the associated read pair information over a region, but this method is not advised for traversing all MUMs or read pairs because the resulting memory access patterns are inefficient.

Information for each MUM is stored in 8 bytes of space. This includes the MUM chromosome (8 bits: up to 255), chromosomal position (32 bits: up to 4.29 billion), MUM length and offset in read (10 bits: up to 1023), plus bits for MUM alignment strand, whether the MUM is on read 1 or read 2, if the MUM is the last MUM in a read pair and if the MUM touches the end of the read.

Read pair information is stored in 8 bytes of space. This includes the index of the first MUM in the read pair (40 bits: up to 1 trillion), the length of each read (10 bits: up to 1023) and bits specifying if the read pair is marked as a duplicate, if the read pair contains MUMs, or if either read is marked as bad by either the sequencer or user software.

Up to 21 bases of unaligned sequence is stored for each read pair in a bases object in 8 bytes of space. If more sequence storage space is needed, the bases object instead contains a 63 bit index to a block in an overflow array encoding all excess sequence in 3 bits per base.

The MUM index for genome ordered access uses 8 bytes per MUM for storing the read pair index (40 bits: up to 1 trillion) and the index of the MUM in the read pair (11 bits: up to 2047), with some space reserved for optional applications. MUMs can be looked up by genome position in O(log N) time and genome position ranges can be traversed in O(R) time, where N is the number of MUMs in the MUMdex file and R is the number of MUMs in the range.

Apart from the flags encoding duplicate and bad read status for a read pair, all MUMdex object access is read-only to prevent inadvertent corruption of data by user code. Pair, MUM and index objects and read sequences are accessed via the MUMdex object either by using integer array indices or by using iterators. MUMdex-related objects are laid out in memory and on disk as simple C++ structures, but access is granted only via overhead-free member functions. The MUMdex object also provides access to a reference object to query various aspects of the genome such as chromosome lengths and sequence.

The format is designed for compactness without employing block compression, so entries can be quickly retrieved in random-access fashion. With a minimum MUM length cutoff of 20, the MUMdex alignment format losslessly compresses real whole genome sequencing reads by about 2.7 times compared to the read sequences stored as ASCII text.

### 3.3 MUMdex Analysis Software and Other Tools

MUM-based analysis begins with one or more MUMdex alignment files. Depending upon project goals the approach taken will differ, but since the MUMdex aligner does not interpret read pairs but simply reports all MUMs found, the alignment format is a generally useful starting point for many types of analysis.

We favor a bridge-based analysis method because it is essentially immune to contamination from base substitution read error for non-SNP candidate types. Bridge-based analysis is performed using the bridges and population_bridges programs (Supplementary Figure 2). The bridges program summarizes information for each bridge observed over all read pairs for a sample and saves the summary by chromosome. The population_bridges program looks for *de novo* candidates and/or events seen in a single family in a region of a chromosome over a population.

Population-based analysis helps to filter false *de novo* candidates resulting from anomalies in the genome reference, common events missed in the parents, and possibly genome regions of great instability. We have prepared a ‘portable’ population database for use by users processing a small number of families who do not have the resources to sequence additional samples. The database contains all bridges seen at least twice in a single individual from a collection of 1020 parents using the hg19 reference, and is included with the Supplementary Materials. Justification for the cutoff of a count of at least two is given in the results.

Other tools distributed in the MUMdex package can be used to convert between formats, facilitate different types of analysis or aid in the examination of MUMdex data:

**bridge_figure:** create an event pdf with explanatory figures

**namepair:** pairs reads in a SAM file by read name

**fastqs_to_sam:** convert fastq files to a name paired SAM file

**count_anchors:** count anchor and reference alleles for a sample

**show_all_counts:** output anchor counts over a population

**anchor_repeatness:** output repetitivity info for bridge anchors

**count_pseudogenes:** find processed pseudogenes in a sample

**denovo_pseudogenes:** find *de novo* candidates over population

**find_bridge:** check samples or families for a specific bridge

**find_microsatellite:** assess microsatellite status for a position

**mumdex2txt:** convert a MUMdex alignment file to text format

**mumdex_sequences:** output the sequence for each read pair

**mumdex2sam:** convert MUMdex alignment file to SAM format

**show_mums:** output MUM information in text format

**show_pairs:** output read pair information in text format

**pair_view:** text view of read pairs to visualize MUM alignments

**bridges2txt:** convert bridges program output to a text format

**karyotype:** create a karyotype figure with event histogram

**population_database:** create a portable population database

**pop2txt:** convert population database between binary and text

## 4 Results

We show that MUMdex software losslessly compresses simulated genomic sequence without error. We characterize performance using real whole genome data from 40 quad (mother, father, proband child and sibling) families of the Simons Simplex Collection (Fischbach *et al*., 2010). We show that *de novo* indel and structural variation candidates can be found using whole genome data from 510 quad families from the same collection.

### 4.1 Verification of MUMdex Correctness Via Simulation

We processed reads from a simulated sample genome containing single base substitutions and structural variation to test the ability of the MUMdex aligner and alignment format to losslessly compress the input. Success for this test is defined as being able to reconstruct the input sequence and read pair mate associations from the MUMdex alignment file while compressing the input sequence better than the gzip (Deutsch 1996) algorithm does.

To produce the simulated sample genome, we first concatenated all chromosomes of the hg19 human reference genome to produce a single sequence. We then introduced base substitutions across the genome with a probability of 0.001 per base. Finally, we introduced breaks between bases in the genome with a probability of 0.001, shuffled the resulting pieces and concatenated them to construct a new sequence. Given the reference genome length of slightly over 3.1 billion bases, this procedure resulted in a simulated sample genome with approximately 3.1 million base substitutions and 3.1 million junctions between sequences not normally adjacent in the human reference genome.

We generated 151 base long paired reads by randomly generating positions in the simulated sample genome uniformly in the range [0, *L*_*G*_ – *L*_*R*_), where *L*_*G*_ is the genome length and *L*_*R*_ is the read length, and extracting the sequence starting at that position. We required that no read sequence selected was identical to any other read sequence to allow definite read identification during correctness testing. 100 million read pairs were generated in this fashion for each of 10 runs for a total of 1 billion read pairs tested.

We ran the MUMdex aligner with a minimum mum length filter of 20 on each group of 100 million read pairs, and every time we were able to reconstruct all input read sequences from the MUMdex alignment file while maintaining the read pair mate associations. On average, the MUMdex alignment file is 4.8 times smaller than the simulated input read sequences stored one sequence per line as ASCII text and 1.4 times smaller than a gzip-compressed version of the same sequence file.

### 4.2 Analysis Speed and Output Size Characteristics

Alignment of 143 bp whole genome read pairs to the hg19 reference and its reverse complement using 12 threads runs at a rate of over 4.5 million read pairs per minute on Xeon E5-2690 processors with sufficient memory and large local Raid-5 SAS storage. Average processing times and storage space requirements for 160 such datasets processed in parallel are shown in Table 1.

**Table 1.** MUMdex time and space. Averages for processing 160 whole genome samples with no MUM length cutoff are shown. The average number of 143 bp read pairs per sample was 439,296,367. The resulting alignment speed of 4.68 million reads per minute and the speed of the merge_mumdex and bridges steps were slower than optimal due to storage device contention from 20 compute nodes processing the samples in parallel. The mummer program used 12 threads per node, bridges used 24 threads per node and the other programs were single threaded.

| Processing Step | Time | Output Size |
| --- | --- | --- |
| mummer | 3.13 hours / sample | 63.214 GB / sample<br>(deleted by next step) |
| merge_mumdex | 3.86 hours / sample | 118.034 GB / sample |
| bridges | 1.55 hours / sample | 47.630 GB / sample |
| population_bridges | 11 hours elapsed | 103 kB total |
| analysis total |  | 165.664 GB / sample |

We compared Bowtie 2 plus SAMtools sort operating as a pipeline with 200GB available for sorting to MUMdex using the same dataset and number of threads (12). We find that Bowtie 2 takes 13 hours versus 7 hours for MUMdex with no MUM length cutoff. MUMdex saves approximately 8 times the number of alignments as Bowtie 2, yet the MUMdex output file is 2/3 the size of the Bowtie 2 BAM file. The MUMdex space advantage is a result of both the reference compression method used and the fact that MUMdex discards basecall quality score information. Operating MUMdex with a MUM length cutoff of 20 greatly reduces the time for the merge_mumdex step and the size of the MUMdex alignment file.

Converting MUMdex output to BAM format results in a BAM file that is about 3% smaller than the MUMdex alignment file. Sequential access of the BAM file is approximately 50% faster than sequential MUMdex access. Random access, lookup by genomic position and especially gathering information for all alignments of a read pair are all much slower for the BAM format than for the MUMdex format. This is because BAM entries are variable sized objects embedded in indexed gzip-compressed blocks that need to be decompressed and linearly searched for each access while the MUMdex format allows direct random access and keeps read pair information together instead of being interspersed as in a BAM file.

### 4.3 Whole Genome *De Novo* Detection

We used MUMdex to analyze whole genome sequence data generated from DNA extracted from whole-blood of 2040 individuals in 510 quads from the Simons Simplex Collection. The libraries had an average fragment length of 362 bases and sequencing was performed in paired end mode to an average depth of coverage of 30. Of the 510 families, 40 were amplified using PCR and had read lengths of 143 bases and the remainder used a ‘PCR-free’ method (Illumina, 2015) and had read lengths of 151 bases. We searched for *de novo* structural variations using stringent conditions.

The stringency of the *de novo* candidate list is defined by setting thresholds for inclusion. We require that each candidate bridge be seen in at least 5 read pairs in a child. In the child, the lengths observed for each bridge MUM must be at least 25 with excess mappability of at least 1 in some read. A bridge with identical characteristics must not be present in the parents of the child, yet the ambient coverage must be at least 10 at both bridge anchor positions in both parents. We also eliminate events as *de novo* if for either bridge anchor in the child a parent has even one MUM with a coincident anchor and that parental MUM has sequence adjacent to the anchor nearly identical to the 10 base adjacent sequence of the child’s anchor.

If a complex event generates multiple bridges, we attempt to retain only the ‘outermost’ of these bridges, meaning that the bridge MUMs must be compatible with the surrounding location. To achieve this, we require 20 or more bases of additional support be seen in at least one mate in the correct orientation for each bridge MUM.

To investigate the effectiveness of the population filter, we allowed the identical bridge characteristics to appear in up to four other families. We call those candidates with no identical bridge seen in any other family the ‘strong’ *de novo* candidates. The other events we call ‘recurrent’. We detected 6278 *de novo* candidates in total, with 4740 being of the strong type. Of the strong candidates, 2376 were found in the proband, 2336 were found in the unaffected sibling and 28 were shared between proband and sibling. Supplementary Figure 3 is a karyogram showing the positions of all anchors of strong candidates. At gross resolution, strong anchors have for the most part a uniform distribution of events over the genome with only a few hotspots and dead zones.

The majority of strong events were short (< 10 bp) deletions, followed by short insertions, larger deletions and insertions, translocations, and even larger insertions and some inversions. Table 2 is a breakdown of observed event types and Supplementary Figure 4 is a histogram of the size distribution of smaller indel events.

**Table 2.**
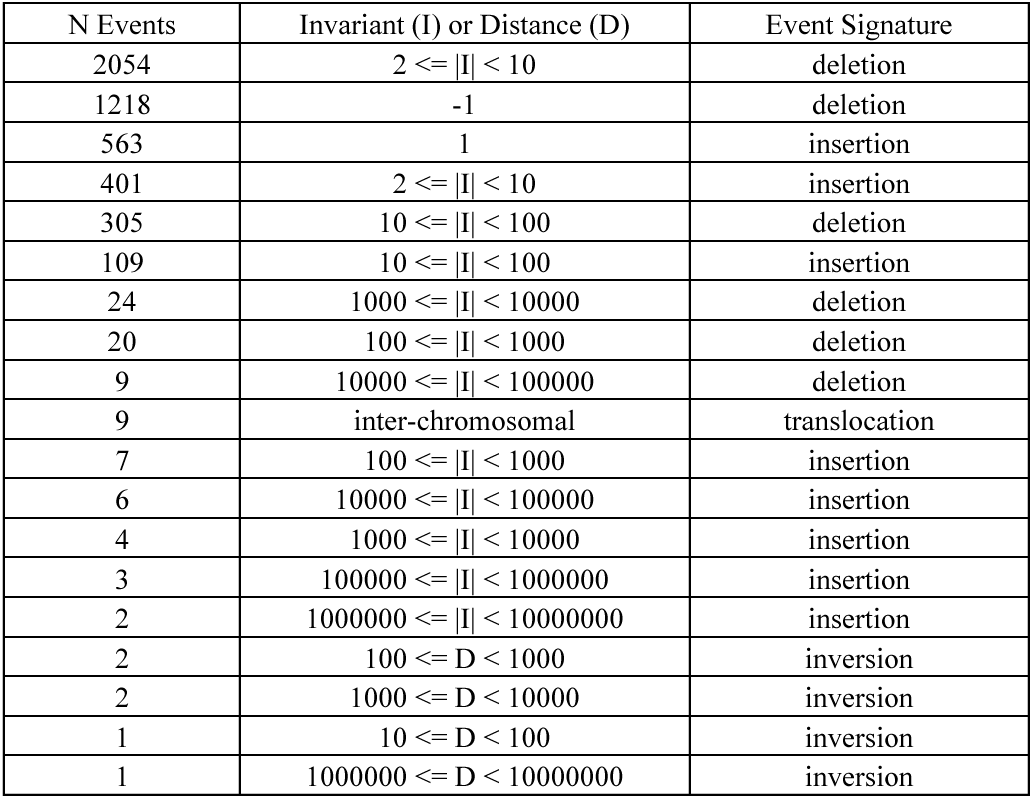
Counts for event type and size range. Event sizes for strong *de novo* candidates are characterized by the invariant (I) for insertions and deletions and by the chromosomal distance (D) between bridge anchors for inversion events.

Events identified as translocations or very large inversions, deletions and insertions should be viewed with skepticism and be investigated in detail. We have examined most of these, and in many cases they arise from a small local de novo event, such as a substitution, which thereby created a spurious MUM, yet the event still passed all filters. Table 1 in the Supplementary Materials lists the properties of each event found in detail.

### 4.4 Properties of recurrent events

Limiting ourselves to strong candidates helps avoid calling a rare variant a *de novo* event when by chance it was missed in the parents. However, the same filter prevents us from identifying true *de novos* that are frequently recurrent. Our full list of *de novos* contains an additional 1538 candidates where the bridge was seen in up to 4 other families.

Figure 4 is a scatter plot of the maximum bridge count seen in other families (which is zero if the event was seen in only one family) versus the candidate bridge count. It demonstrates that in the vast number of cases, the bridge was either not seen or was seen at a maximum count of 1 in some other family. Events seen at a count of more than 1 in some other family show the expected roughly linear relationship between the candidate bridge count and the maximum other family bridge count. Events seen with a count greater than one in some other family also appear to transmit roughly as expected (Supplementary Figure 5), strongly suggesting that they are not somatic mutations or artifacts of library preparation or sequencing platform. By contrast, almost all events seen at a maximum count of one in other families are not transmitted, and thus are either somatic mutation or an artifact of preparation or platform in those families.

**Figure 4.**
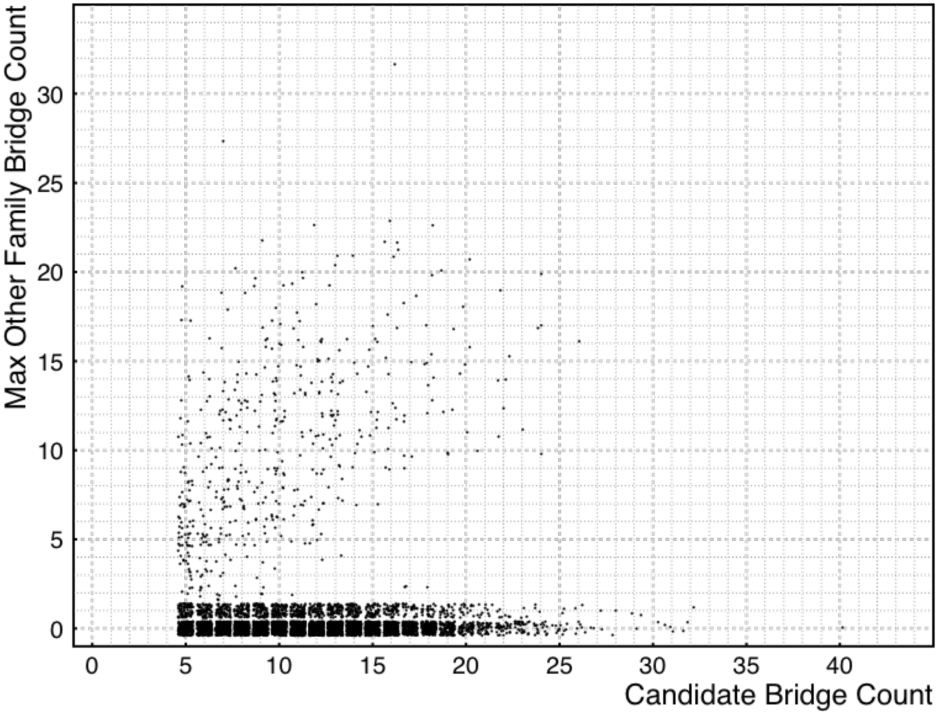
Maximum bridge counts. For every candidate *de novo* event (seen in a child, not its parents and fewer than five other families) we plot its bridge count on the X-axis vs. the maximum bridge count observed in any non-candidate family on the Y-axis. Jitter has been added to the counts to help separate overlapping points.

The group of recurrent candidates (those seen with high bridge counts in other families) have significantly different properties than the strong candidates, and we believe these properties may have been the cause of much of the recurrence. As shown in Figure 5 Panel A, the offset between bridge anchor positions tends to be negative for events seen in more than one family. The effect is stronger still for events that are transmitted (Panel B). A negative offset means that the bridge MUMs have high overlap, which is expected for repeat expansion and contraction events and for non-allelic homologous recombination events. These observations are consistent with finding long repeats more frequently at the bridge anchor coordinates of recurrent *de novos* (see Supplementary Figure 7).

**Figure 5.**
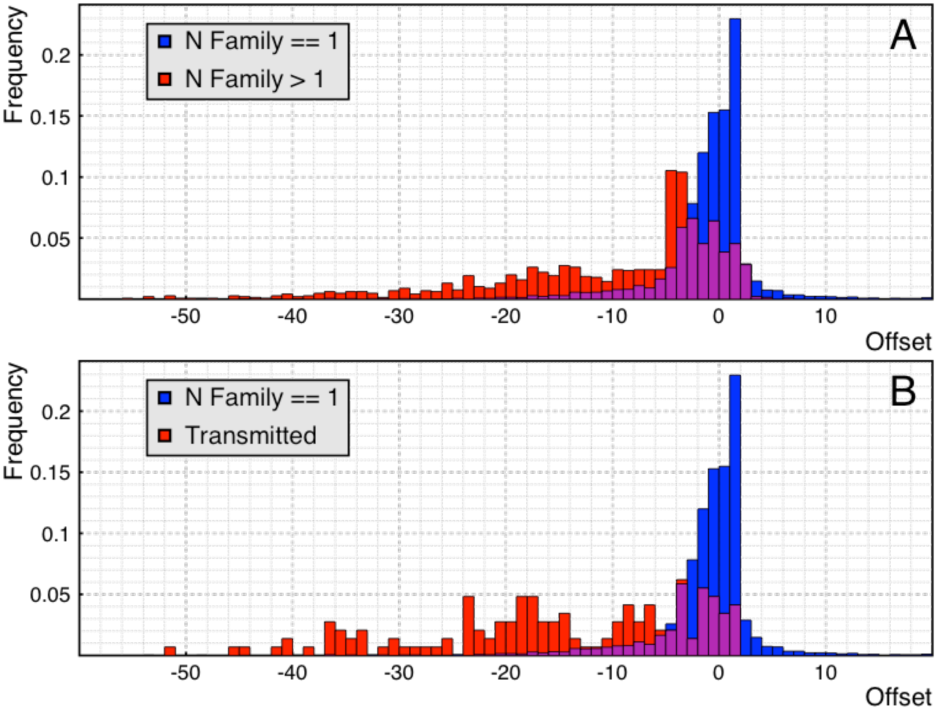
Offsets for recurrent and non-recurrent candidates. Negative offset values indicate that the bridge MUMs overlap each other. In blue in both panels we see a normalized histogram of anchor read offsets of strong *de novo* candidates. **Panel A** shows a normalized histogram of anchor read offsets for candidates seen in from two to five families (red). The area in common between the two histograms is displayed in purple. **Panel B** shows a normalized histogram of anchor read offset for candidates seen in at least two members, each at a count of at least two, in another family (red), a proxy for transmission.

The interpretation of Figure 4 provides the justification for the bridge inclusion criteria in our portable population database, where we only include a bridge if seen at least twice in at least one individual. A scatter plot of invariant vs position for indels from the portable database is shown in Supplementary Figure 6 for a typical one megabase region. Clearly, the aggregate events in the human population show a highly structured distribution, mostly reflecting the repeat structure in the genome.

The ~250 to 700 base long deletion events in Supplementary Figure 6 seen ubiquitously in the population filter demonstrate its utility. As discussed in the figure caption, these apparent ‘deletions’ are predominantly a matter of library preparation, present at ~25 fold the incidence in the ‘PCR-free’ libraries than in the conventionally prepared libraries. They are recurrent artifacts caused by the PCR-free sequencing method, and the recurrence filter effectively rejects them as *de novo* events.

## 5 Discussion

Specific gross changes in genome structure and copy number have long been known to be the cause of various syndromes (Lejeune *et al*., 1963; Lupski *et al.*, 1991) and play a major role in cancer (Lugo *et al.*, 1990; Slamon *et al.,* 1989). Structural variation in the human genome will surely play a large role in our ultimate understanding of evolution, heredity and disease. Yet, the reference genome sequence assembly is weakest where variation of this type is greatest, and its analysis is far from complete. Analytic tools are critically needed now that thousands of deep whole genome sequences are becoming available. We offer MUMdex as one such tool.

Several key features distinguish MUMdex from other analysis methods. MUMdex uses only unique matches to a reference genome assembly. MUMdex saves all MUMs and unaligned sequence to create a general-purpose sequencing run index, delays the interpretation of reads until all information is available, characterizes events by their bridge coordinates and invariants and, additionally, uses a population to help filter events. The population database aspect of MUMdex analysis helps avoid false events due to an imperfect reference genome and fully utilizes the large family-based WGS datasets available to us from the SSC (Fischbach *et al.,* 2010).

The extensive use of maximum unique matches (MUMs) is a choice that places MUMdex at an extreme point in a space of computational tradeoffs. We contrast it to other methods with anchor finding techniques that permit multiple or mutated matches to the genome, which are limited to local genome searches (and hence the discovery of only smallscale structural variation) because otherwise there are too many possibilities. What is lost for MUMdex in the tradeoff is the ability to identify events in duplicated regions or in regions highly divergent from the reference assembly. Care must also be taken when interpreting MUMdex events, as the true nature of an event may not be obvious. For instance, an event with the signature of a translocation may actually be a SNP that induced a spurious MUM to a distant but similar sequence.

In this manuscript we have presented MUMdex software and described the principles behind our analysis, to make it available for use by the community at large. We have used MUMdex to explore *de novo* structural variation in a large population. We have made a ‘portable’ database of structural variants and recurrent spurious events available for anyone seeking to use our software but who does not have access to large family databases.

While we are working on validating these preliminary results in the context of autism studies, the broad outline of what we have found so far is likely to survive any validations. There is no bias to discovery of ‘local’ events by MUMdex, yet most events are small deletions and insertions, with frequency decreasing as event size increases. We observe that recurrent events are more likely to be found in repetitive regions, and these regions are likely to be inherently unstable.

In addition to extensive validation, in our future work we will refine criteria for filtering *de novo* events, explore regions of genome instability, and craft ‘interpretative’ tools, such as the bridge_figure program mentioned in Software (see Supplementary Figure 8), that allow the manual inspection of events.

## Acknowledgements

We thank all the families at the participating Simons Simplex Collection (SSC) sites, as well as the principal investigators (A. L. Beaudet, R. Bernier, J. Constantino, E. H. Cook Jr, E. Fombonne, D. Geschwind, D. E. Grice, A. Klin, D. H. Ledbetter, C. Lord, C. L. Martin, D. M. Martin, R. Maxim, J. Miles, O. Ousley, B. Peterson, J. Piggot, C. Saulnier, M. W. State, W. Stone, J. S. Sutcliffe, C. A. Walsh and E. Wijsman) and the coordinators and staff at the SSC sites for the recruitment and comprehensive assessment of simplex families; and the SFARI staff for facilitating access to the SSC.

## References

Cerf V. (1969) ASCII format for Network Interchange, Network Working Group RFC, 20, Retrieved from http://www.faqs.org/rfcs/rfc20.html

Chen K, Wallis J, McLellan M, Larson D, Kalicki J, Pohl C, McGrath S, Wendl M, Zhang Q, Locke D, Shi X, Fulton R, Ley T, Wilson R, Ding L, Mardis E (2009) Breakdancer: an algorithm for high-resolution mapping of genomic structural variation, Nature Methods, 6 (9), 677–684

Cock P, Fields C, Goto N, Heuer M, Rice P. (2010) The Sanger FASTQ file format for sequences with quality scores, and the Solexa/Illumina FASTQ variants, Nucleic Acids Research, 6, 1767–1771

Deutsch P. (1996) GZIP file format specification version 4.3, Network Working Group RFC, 1952, Retrieved from https://tools.ietf.org/html/rfc1952

Fischbach GD, Lord C. (2010) The Simons Simplex Collection: a resource for identification of autism genetic risk factors. Neuron, 68 (2), 192–195

Illumina. (2015) TruSeq DNA PCR-Free Library Prep Reference Guide. Part # 15036187 Rev. D Catalog # FC-121-9006DOC

International Organization for Standardization (ISO). (2014) ISO International Standard ISO/IEC 14882:2014(E) - Programming Language C++. Retrieved from http://isocpp.org/std/the–standard

Karakoc E, Alkan C, O’Roak B, Dennis M, Vives L, Mark E, Rieder M, Nickerson D, Eichler E. (2012) Detection of structural variants and indels within exome data. Nature Methods, 9 (2), 176–180

Kasai T, Lee G, Arimura H, Arikawa S, Park K. (2001) Linear-Time Longest-Common-Prefix Computation in Suffix Arrays and Its Applications, Proceedings of the 12th Annual Symposium on Combinatorial Pattern Matching, Lecture Notes in Computer Science 181–192

Khan Z, Bloom JS, Kruglyak L, Singh M. (2009) A practical algorithm for finding maximal exact matches in large sequence datasets using sparse suffix arrays, Bioinformatics, 25 (13), 1609–1616

Langmead, B, Salzberg S. (2012) Fast gapped-read alignment with Bowtie 2, Nature Methods, 9 (4), 357–359

Lejeune J, Lafourcade J, Berger R, Vialatte J, Boeswillwald M, Seringe P, Turpin R. (1963). 3 Cases of partial deletion of the short arm of chromosome 5. C. R. Hebd. Seances Acad. Sci. (in French). 257: 3098–3102.

Li H, Durbin R. (2009) Fast and accurate short read alignment with Burrows-Wheeler transform, Bioinformatics, 25 (14), 1754–1760

Li H, Handsaker B, Wysoker A, Fennell T, Ruan J, Homer N, Marth G, Abecasis G, Durbin R, 1000 Genome Project Data Processing Subgroup. (2009) The Sequence Alignment/Map format and SAMtools, Bioinformatics, 25 (16), 2078–2079

Lindberg M, Hall I, Quinlan A. (2015) Population-based structural variation discovery with Hydra-Multi, Bioinformatics, 31 (8), 1286–1289

Lugo TG, Pendergast AM, Muller AJ, Witte ON. (1990) Tyrosine kinase activity and transformation potency of bcr-abl oncogene products. Science. 247:1079–1082.

Lupski JR, De Oca-Luna RM, Slaugenhaupt S, Pentao L, Guzzetta V, Trask BJ, Saucedo-Cardenas O, Barker DF, Killian JM, Garcia CA, Chakravarti A, Patel PI. (1991) DNA duplication associated with Charcot-Marie-Tooth disease type 1A. Cell. 66: 219–232.

Manber U, Myers G. (1993) Suffix Arrays: A New Method for On-Line String Searches, SIAM Journal on Computing 22 (5) 935

McKinney EH. (1966) Generalized Birthday Problem, American Mathematical Monthly, 73, 385–387

Slamon DJ, Godolphin W, Jones LA, Holt JA, Wong SG, Keith DE, Levin W, Stuart S, Udove J, Ullrich A, Press M. (1989) Studies of the HER-2/neu protooncogene in human breast and ovarian cancer. Science 244: 707–712.

Ye K, Schulz M, Long Q, Apweiler R, Ning Z. (2009) Pindel: a pattern growth approach to detect break points of large deletions and medium sized insertions from paired-end short reads, Bioinformatics, 25 (21), 2865–2871

